# Multilocus sequence analysis, a rapid and accurate identification tool for taxonomic classification, evolutionary relationship and population biology of the genus *Shewanella*

**DOI:** 10.1101/512624

**Authors:** Yujie Fang, Yonglu Wang, Zongdong Liu, Hang Dai, Hongyan Cai, Zhenpeng Li, Zongjun Du, Xin Wang, Huaiqi Jing, Qiang Wei, Biao Kan, Duochun Wang

**Author notes:** Correspondence Author: Duochun Wang,. Changbai Road 155, Changping, Beijing, 102206, China.

## Abstract

The genus *Shewanella* comprises a group of marine-dwelling species with worldwide distribution. Several species are regarded as causative agents of food spoilage and opportunistic pathogens of human diseases. In this study, a standard multilocus sequence analysis (MLSA) based on six protein-coding genes (*gyrA, gyrB, infB, recN, rpoA* and *topA*) was established as a rapid and accurate identification tool in fifty-nine type *Shewanella* strains. This method yielded sufficient resolving power in regard to enough informative sites, adequate sequence divergences and distinct interspecies branches. The stability of phylogenetic topology was supported by high bootstrap values and concordance with different methods. The reliability of the MLSA scheme was further validated by identical phylogenies and high correlations of genomes. The MLSA approach provided a robust system to exhibit evolutionary relationships in the *Shewanella* genus. The split network tree proposed twelve distinct monophyletic clades with identical G+C contents and high genetic similarities. Eighty-six tested strains were investigated to explore the population biology of the *Shewanella* genus in China. The most prevalent *Shewanella* species were *Shewanella algae, Shewanella xiamenensis, Shewanella chilikensis, Shewanella indica, Shewanella seohaensis* and *Shewanella carassii*. The strains frequently isolated from clinical and food samples highlighted the importance of increasing the surveillance of *Shewanella* species. Combined with the genetic, genomic and phenotypic analyses, *Shewanella upenei* should be considered a synonym of *S. algae*, and *Shewanella pacifica* should be reclassified as a synonym of *Shewanella japonica*.

**IMPORTANCE:** The MLSA scheme based on six HKGs (*gyrA, gyrB, infB, recN, rpoA* and *topA*) is well established as a reliable tool for taxonomic, evolutionary and epidemiological analyses of the genus *Shewanella* in this study. The standard MLSA method allows researchers to make rapid, economical and precise identification of *Shewanella* strains. The robust phylogenetic network of MLSA provides profound insight into the evolutionary structure of the genus *Shewanella*. The population genetics of *Shewanella* species determined by the MLSA approach plays a pivotal role in clinical diagnosis and routine monitoring. Further studies on remaining species and genomic analysis will enhance a more comprehensive understanding of the microbial systematics, phylogenetic relationships and ecological status of the genus *Shewanella*.

---

The genus *Shewanella*, first described by MacDonell & Colwell, belongs to the family *Shewanellaceae* as a sole genus (1). The members of this genus are gram-negative, facultatively anaerobic, oxidase-positive and motile bacteria (2–4). At the time of writing, there are more than sixty recognized species in the genus of *Shewanella* (http://www.bacterio.net/shewanella.html). The majority of *Shewanella* species inhabit a wide range of environments, including free-living in oceans (5–8). The genus *Shewanella* plays a critical role in bioremediation (9), and certain strains have been used in bioelectrical systems (10, 11). In addition, multiple *Shewanella* species are frequently yielded from consumable products as spoilage bacteria and clinical specimens as opportunistic pathogens (12–14).

To date, polyphasic approaches are performed to assign the phylogenetic placement and taxonomic classification of *Shewanella* species. Commercial biochemical systems, such as Vitek and API, are available for species identification in clinical laboratories. However, only two species, namely, *S. algae* and *Shewanella putrefaciens*, have been recorded in the database (12, 13). Phylogenetic analysis based on the 16S rRNA gene as a molecular marker was utilized to yield an evolutionary relationship for taxa (15). The disadvantage of the application of the 16S rRNA gene was the low resolving power to discriminate closely related species due to their high sequence similarities (16). Recently, a more rapidly evolving housekeeping gene (HKG) of *gyrB* was selected as an alternative phylogenetic indicator for *Shewanella* species classification (17–20). Nevertheless, the quality of sequences submitted in public databases is poor (20–22). The genome-wide parameters, consisting of *in silico* DNA-DNA hybridization (isDDH) (23) and average nucleotide identity (ANI) (24), take the place of the wet-lab DDH to unravel bacterial systematics. However, the process of genome sequencing is expensive and time-consuming; meanwhile, limited genomes of type *Shewanella* species are available in public databases. These conditions make this approach impractical in clinical and daily investigations for rapid and efficient identification.

The effective MLSA scheme has been applied to increasing prokaryotic taxa, for instance, the genera of *Virbio* (25, 26), *Aeromonas* (27), *Enterobacter* (28), *Treponema* (29) and *Halomonadaceae* (30). Nevertheless, rare information is delineated among the genus *Shewanella*. Hence, in this study, we established a reliable MLSA method to classify *Shewanella* species by assessing the nucleotide sequences and phylogenies of six individual and concatenated HKGs (*gyrA, gyrB, infB, recN, rpoA* and *topA*) in almost sixty type *Shewanella* strains. The phylogenetic framework of concatenated sequences provided a significant understanding of the evolutionary relationship in the genus *Shewanella* on the basis of multiple distinct taxonomic clades. The MLSA scheme was further utilized to determine the population biology of eighty-six tested strains collected in China.

## MATERIALS AND METHODS

### Bacterial strains and culture conditions

A total of 145 *Shewanella* strains were involved in this study. Forty-two type strains were collected from the China General Microbiological Culture Collection Center (CGMCC), the German Collection of Microorganisms and Cell Cultures (Deutsche Sammlung von Mikroorganismen und Zellkulturen, DSMZ), the Japan Collection of Microorganisms (JCM), Korean Collection for Type Cultures (KCTC), LMG Bacteria Collection (LMG) and Marine Culture Collection of China (MCCC); the other seventeen type strains with complete genomes that were available in GenBank were used for sequence analyses; eighty-six tested strains were isolated from patients (n = 44), food (n = 35) and environments (n = 7) in four provinces (Anhui, Hainan, Liaoning and Shandong) of China from 2007 to 2016. Detailed type strain information is listed in Table S1. The forty-two type *Shewanella* strains were incubated at suitable conditions following the protocols of culture collection. The tested strains were cultured on Marine Agar 2216 (BD, Difco) at 35 °C for 18 h.

### DNA extraction, gene selection and primer design

Genomic DNA from *Shewanella* strains was extracted with a genomic DNA extraction kit (TaKaRa, Dalian, China) following the manufacturer’s instructions. The 16S rRNA gene of tested strains was amplified and sequenced with two universal primers (27F and 1492R) described previously (31). Six HKGs (*gyrA, gyrB, infB, recN, rpoA* and *topA*) were chosen for the MLSA scheme. The degenerate primers of HKGs for PCR amplification, except the *gyrB* gene referring to Yamamoto & Harayama (32), were designed from genome sequences of type *Shewanella* strains in the GenBank database (Table S1) to accommodate a wide taxonomic scope. The nondegenerate primers on the 5’ region for sequencing are underlined in Table S2.

### PCR amplification and sequencing

Amplification reactions for six HKGs were performed in a total volume of 25 μl, containing 12.5 μl 2× Es Taq MasterMix (Cwbiotech, China), 2 μl each forward and reverse primer (10 μM), 1.5 μl template DNA (10-30 ng/μl) and 7 μl ultrapure water using SensoQuest LabCycler. The PCR mixture was subjected to denaturation at 94 °C for 10 min, followed by 35 cycles of denaturation at 94 °C for 30 s, annealing at 54-60 °C for 30 s and extension at 72 °C for 60 s, with a final extension step at 72 °C for 10 min. More detailed information on annealing temperatures is listed in Table S2. PCR amplicons were verified by electrophoresis on 1 % agarose TBE gels at 220 V for 15 min, stained with GoldView (Solarbio, China), and visualized on a UV transilluminator with a clear single band at the expected length. The amplified products were purified and sequenced with the ABI 3730xl platform by Tsingke Corporation (Beijing) using the corresponding sequencing primers (Table S2).

### Analysis of nucleotide diversity

The sequences of 16S rRNA, *gyrA, gyrB, infB, recN, rpoA* and *topA* genes used for MLSA were trimmed to positions 56-1455, 247-744, 337-1446, 1519-2181, 565-1200, 139-756 and 106-768, respectively, corresponding to *E. coli* numbering (33). The evolutionary distances and sequence similarities of the 16S rRNA gene, individual and concatenated HKGs were calculated using MEGA 6.06 (34) with Kimura’s 2-parameter model. The parsimony informative sites and *Ka/Ks* ratios (*Ka*: the number of nonsynonymous substitutions per nonsynonymous site, *Ks*: the number of synonymous substitutions per synonymous site) were analyzed with DnaSP 6.0 (35).

### Phylogenetic analysis

The nucleotide sequences were aligned using MEGA 6.06 (34). The phylogenetic trees of the 16S rRNA gene and the individual and concatenated sequences of six HKGs (*gyrA, gyrB, infB, recN, rpoA* and *topA*) were constructed by neighbor-joining and maximum-likelihood methods with MEGA 6.06. The model selected was Kimura’s two-parameter with the pairwise-deletion option. The robustness of tree topologies was evaluated with 1000 bootstrap replications, and values greater than 70 % are shown at nodes of branches. The split network tree of MLSA was performed by SplitsTree 4.14.4 using the Jukes-Cantor correlation.

### Genomic relatedness

Twenty-eight type *Shewanella* strains with complete genomes available in GenBank (Table S1) were involved to investigate the concordance and correlation between MLSA and genomes. Core genes of genomic sequences identified by OrthoMCL 2.0.9 were concatenated to construct the phylogenetic tree. The *is*DDH results were measured by the Genome-to-Genome Distance Calculator (GGDC) (http://ggdc.dsmz.de/). The values of ANI were estimated by the web-based platform EZBioCloud (http://www.ezbiocloud.net/tools/ani) with the OrthoANIu algorithm. The correlation between the *is*DDH results and MLSA similarities was simulated using MATLAB R2016a (Math Works Inc., USA) with nonlinear interpolation analysis.

### Phenotypic characteristics

Further phenotypic tests were performed among controversial *Shewanella* species whose *is*DDH values were greater than the species threshold. The type strains of species were examined in parallel under suitable conditions. Physiological and biochemical traits were determined by commercial strips, including API 20E and API 20NE (BioMérieux, France), in agreement with the standard manufacturer’s instructions.

## RESULTS

### Individual gene analysis

In this study, sequence diversity and phylogenetic analysis of fifty-nine type strains (Table S3) were performed to assess the interspecies taxonomy among the genus *Shewanella*. The results of sequence diversity for the 16S rRNA gene are shown in Table 1. The high occurrences of greater than 98.65 % interspecies similarity in the 16S rRNA gene implied the low resolution to distinguish *Shewanella* species. The low bootstrap values indicated the unstable topology in the phylogenetic tree, and close evolutionary branches were discovered (Fig. S1). Among the six HKG analyses, greater values of parsimony informative sites and nucleotide diversity were obtained (Table 1). In addition, the phylogenetic trees of all HKGs demonstrated more distinct branches and greater bootstrap values in contrast with the 16S rRNA gene (Fig. S1). However, it was not sufficient to differentiate all members of the genus *Shewanella*. Lower bootstrap values for the outer branches and discordance in the partial topology of six HKGs were still observed.

**Table 1.** Nucleotide sequence diversity of fifty-nine *Shewanella* type strains.

| Locus | Length (bp) | Parsimony<br>informative sites |  | Nucleotide<br>diversity, Pi | Similarities (%) |  | Ka/Ks |
| --- | --- | --- | --- | --- | --- | --- | --- |
|  |  | No. | % |  | Range | Mean |  |
| 16S rRNA | 1434 | 148 | 10.3 | 0.043 | 89.8-100 | 95.0 | NA |
| *gyrA* | 498 | 229 | 46.0 | 0.223 | 68.3-100 | 77.7 | 0.117 |
| *gyrB* | 1110-1119 | 492 | 44.0 | 0.194 | 73.2-99.9 | 80.8 | 0.089 |
| *infB* | 663 | 289 | 43.6 | 0.193 | 73.5-100 | 80.7 | 0.105 |
| *recN* | 633-636 | 457 | 71.9 | 0.360 | 52.2-99.8 | 64.0 | 0.275 |
| *rpoA* | 615 | 221 | 35.9 | 0.125 | 79.7-100 | 87.5 | 0.052 |
| *topA* | 657-660 | 358 | 54.2 | 0.264 | 65.9-100 | 73.5 | 0.168 |
| MLSA | 4176-4191 | 2046 | 48.8 | 0.223 | 71.1-99.9 | 77.7 | 0.143 |

### Multilocus sequence analysis (MLSA)

The concatenated sequences of protein-coding genes for fifty-nine type *Shewanella* strains comprised 2046 (48.8 %) parsimony informative sites with a nucleotide diversity value of 0.223 (Table 1). The analysis of sequences indicated that the MLSA scheme possessed an appropriate resolution and balanced the divergent evolutionary rates of six HKGs. The neighbor-joining phylogenetic tree based on concatenated alignment showed independent branches for interspecies, except for two sets of species, species *S. algae*-*S. haliotis*-*S. upenei* and *S. japonica*-*S. pacifica* (Fig. 1). Those five species were likely to be misclassified, and more approaches were needed to perform the identification. The branches to discriminate *Shewanella* species were supported by high bootstrap values, except for species *S. algicola-S. inventionis* and *S. carassii*. Bootstrap results indicated that the taxonomic groups involving those three species shared close evolutionary relationships. The phylogenetic tree of concatenated sequences was also reconstructed by the maximum-likelihood algorithm (Fig. S2). Almost the same topology was obtained, and the only exception was the location of species *S. carassii*, which was supported with relatively low bootstrap values as described above.

**Figure 1.**
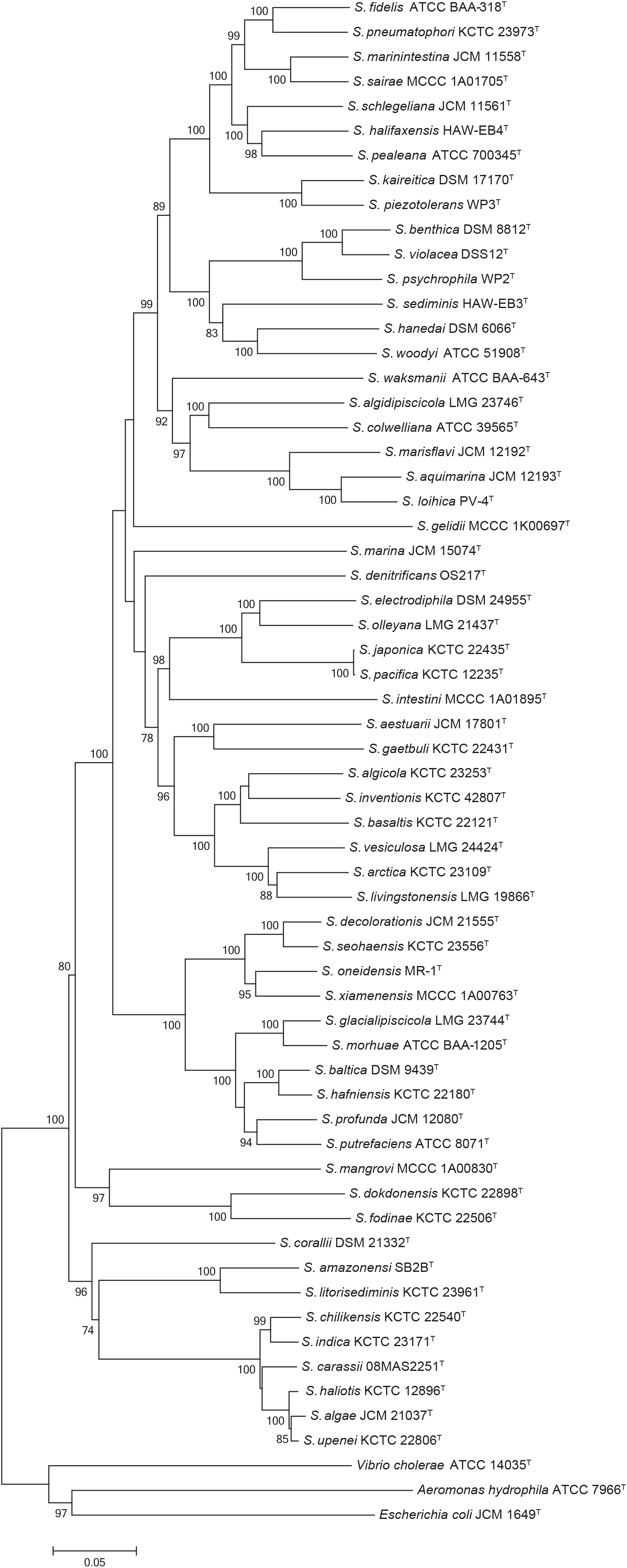
Phylogenetic tree reconstructed by the neighbor-joining method based on concatenated six gene sequences (*gyrA, gyrB, infB, recN, rpoA* and *topA*, 4191 bp) of fifty-nine *Shewanella* type strains. The robustness of tree topologies was evaluated with 1000 bootstrap replications, and values greater than 70 % were shown at nodes of branches. The scale bar indicates substitutions per site. The type strains of *Aeromonas hydrophila* ATCC 7966^T^, *Escherichia coli* JCM 1649^T^ and *Vibrio cholerae* ATCC 14035^T^ served as outgroups.

### Comparative analysis between MLSA and genomes

To further validate the reliability of MLSA, a whole-genome-based phylogenetic tree was constructed, and correlation analysis was performed among twenty-eight type strains whose genomes were publicly available. The phylogeny of MLSA yielded a similar topology to that of core genes, and only a slight difference was observed in the position of species *S. carassii* (Fig. S3). Similarities of the MLSA and *is*DDH were calculated and are shown in Table S4. The *is*DDH values among distant species were concentrated at 20 %. The *is*DDH values were highly correlated with the MLSA similarities (R^2^ = 0.9887) in closely related *Shewanella* species (Fig. 2). Based on the simulative equation of y = 90.01*exp(0.001112*x)-431.3*exp(−0.1927*x), the 70 % *is*DDH value was equivalent to the 97.3 % MLSA similarity, which could serve as a species boundary in the genus *Shewanella*.

**Figure 2.**
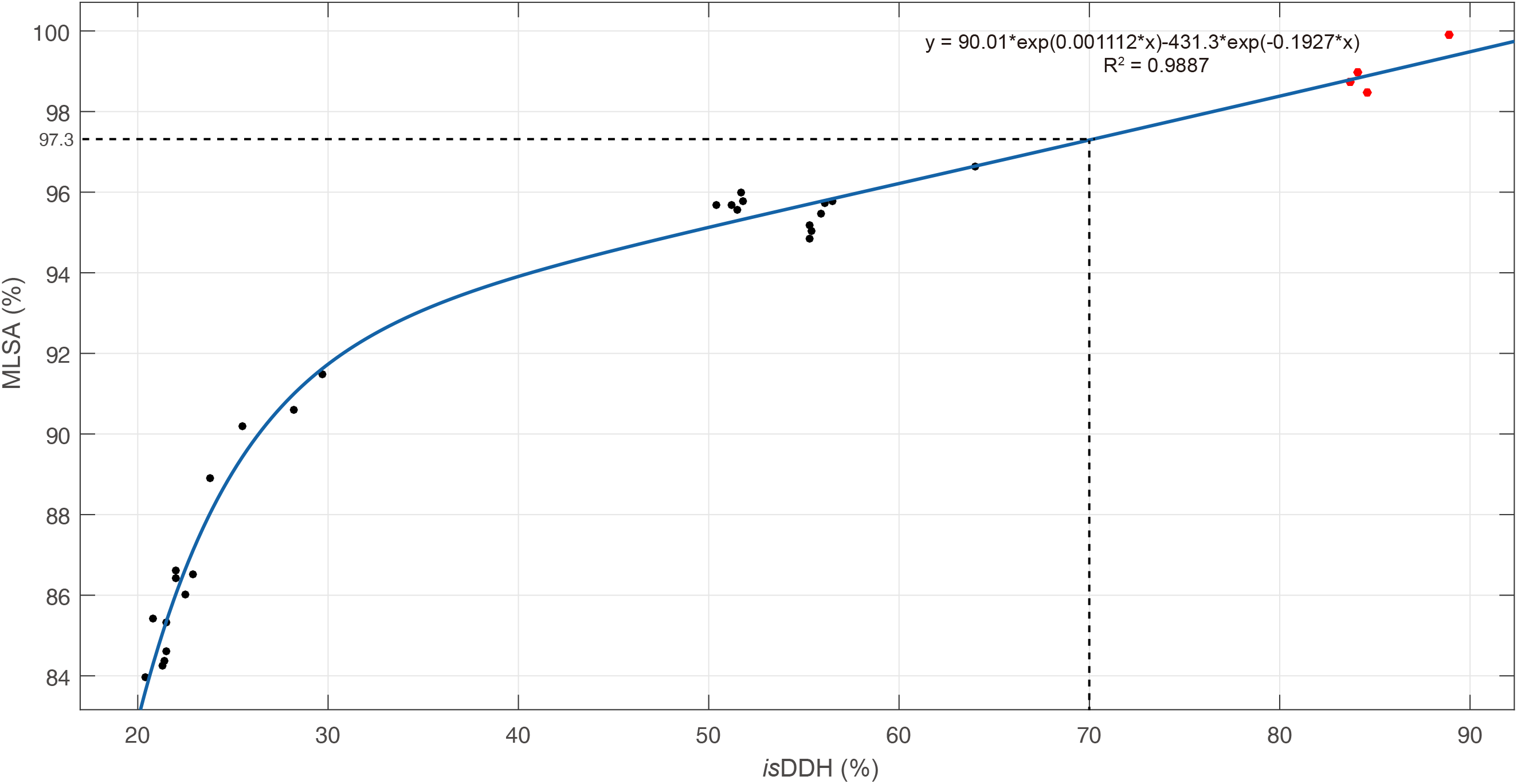
The correlation analysis between similarities of *is*DDH and MLSA for the genus *Shewanella*. The vertical line indicates a 70 % *is*DDH threshold, and the horizontal line indicates the corresponding 97.3 % MLSA similarity. The four points greater than the species boundary are marked in red.

Nevertheless, greater than 97.3 % concatenated sequence similarities were observed among two sets of species, i.e., *S. algae-S. haliotis-S. upenei* and *S. japonica*-*S. pacifica*. The corresponding *is*DDH results between those groups of species were in the range of 83.7-88.9 %, which exceeded the 70 % species threshold (Fig. 2). The further pairwise ANI results between type strains of *S. algae, S. haliotis* and *S. upenei* were 98.2, 98.1 and 98.2 %, respectively, and the value of that between species *S. japonica* and *S. pacifica* was 98.8 %. All ANI values were greater than the boundary of 95 % for species delineation. The genomic analysis based on *is*DDH and ANI provided compelling evidence for correct taxonomic position, indicating that *S. algae, S. haliotis* and *S. upenei* were the same species and *S. pacifica* belonged to the species *S. japonica*. Additional phenotypic characteristics were detected among these five strains (Table 2). Minor differences in biochemical results were obtained between species *S. algae, S. haliotis* and *S. upenei*. The phenotypic discrepancies between species *S. japonica* and *S. pacifica* were discovered in growth conditions and the assimilation of *N*-acetyl-glucosamine. These results confirmed the conclusion of a recent report that identified *S. haliotis* as a synonym of *S. algae* according to whole-genome sequencing. Considering the genetic, genomic and phenotypic characteristics, species *S. upenei* reported by Kim *et al*. 2011 should be regarded as a later heterotypic synonym of *S. algae* Simidu *et al*. 1990, meanwhile, *S. pacifica* Ivanova *et al*. 2004 should be reclassified as a later heterotypic synonym of *S. japonica* Ivanova *et al*. 2001.

**Table 2.**
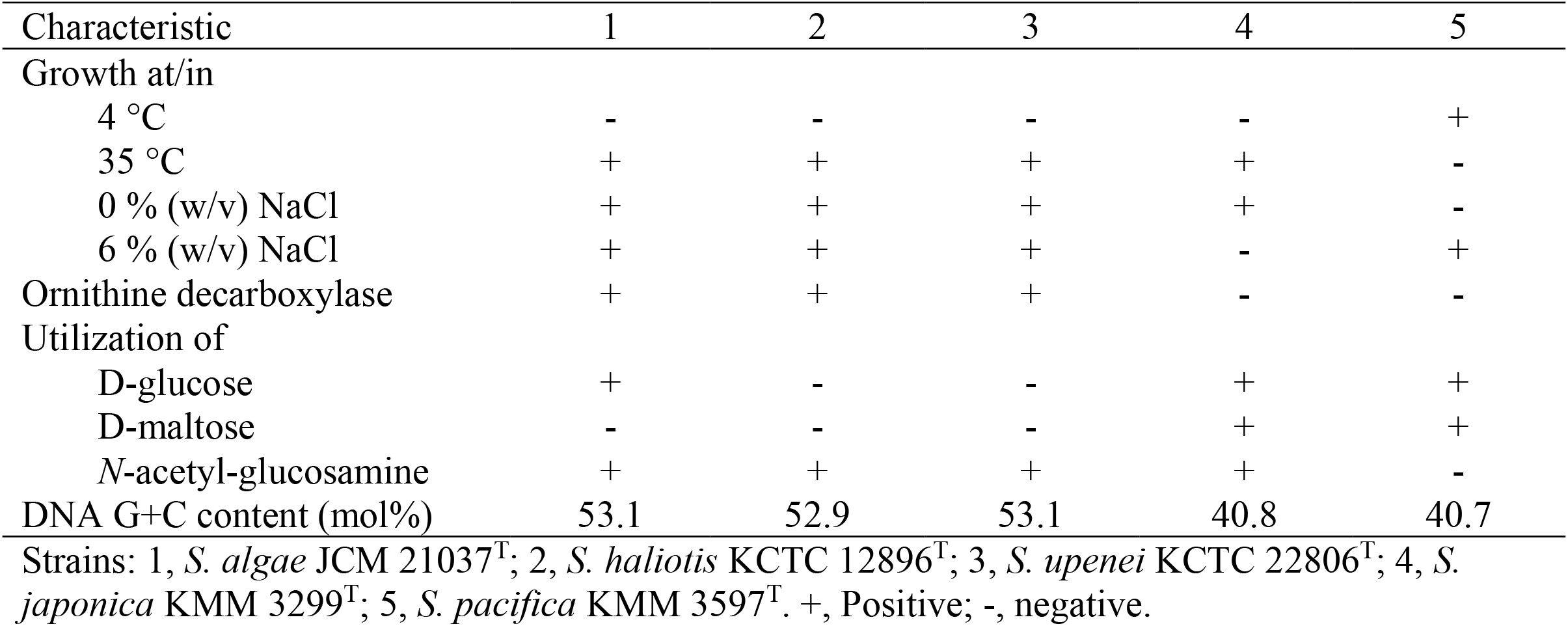
Distinctive phenotypic characteristics between five controversial *Shewanella* strains.

### Distinct taxonomic clades

Given the results of sequence diversity, topological stability and concordance with genomes, the MLSA scheme of six protein-coding genes (*gyrA, gyrB, infB, recN, rpoA* and *topA*) was validated for taxonomic and evolutionary analysis among the *Shewanella* genus. The concatenated sequences for fifty-six species after emendation were subjected to construct the split network tree to explore evolutionary relationships among taxa (Fig. 3). Twelve distinct monophyletic clades were identified, i.e., Algae, Amazonensis, Aquimarina, Benthica, Colwelliana, Fodinae, Gaetbuli, Hanedai, Japonica, Livingstonensis, Pealeana and Putrefaciens clades (Table 3). The *Shewanella* species within the same clade shared <4 mol% GC variation and >84 % MLSA concatenated similarity. There are eight orphan *Shewanella* species, namely, *S. corallii, S. denitrificans, S. gelidii, S. intestini, S. mangrovi, S. marina, S. sediminis* and *S. waksmanii*, which form a distinct branch clearly separated from all taxonomic clades in the phylogenetic network, except for *S. sediminis*. Species *S. sediminis* harbored a far evolutionary distance similar to both Hanedai and Benthica clades and was located on the boundary of clade differentiation. Combined with the ambiguous relationships between species *S. sediminis* and clades Hanedai and Benthica in a single HKG phylogenetic tree, *S. sediminis* was considered an orphan species. Twelve evolutionary clades were always maintained in phylogenetic trees of individual and concatenated HKGs. There were only slight differences observed, i.e., species *S. woodyi-S. hanedai* in *gyrB*, species *S. colwelliana*-*S. algidipiscicola* and *S. gaetbuli*-*S. aestuarii* in *infB*, and species *S. algidipiscicola*-*S. colwelliana* in *topA*, which were positioned closely but did not group within one clade in phylogenies.

**Figure 3.**
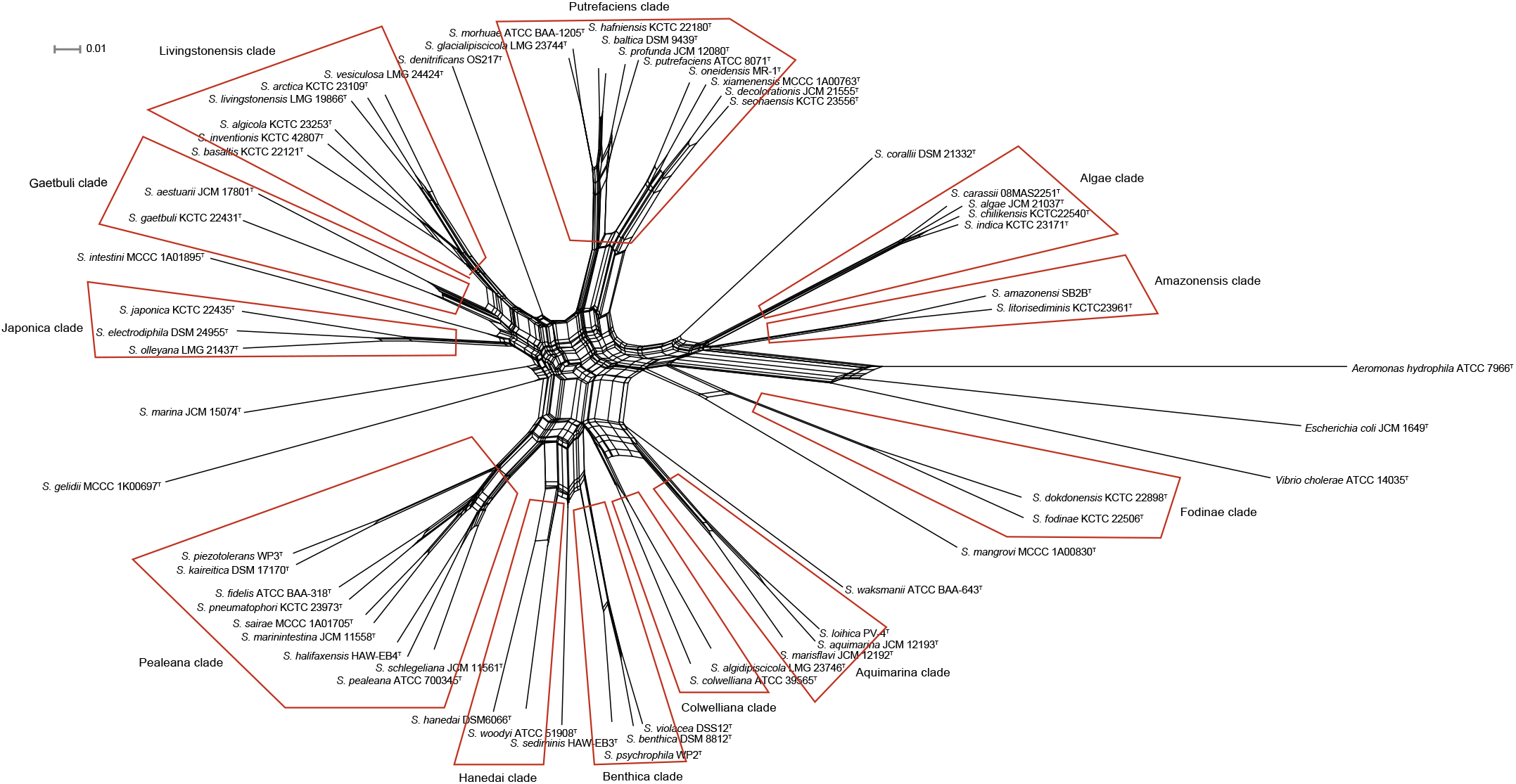
Concatenated split network tree based on six gene loci. The *gyrA, gyrB, infB, recN, rpoA* and *topA* gene sequences from fifty-six validated *Shewanella* species were concatenated and reconstructed using the SplitsTree 4 program. Twelve distinct clades were identified and indicated by a red line.

**Table 3.** G+C content and MLSA concatenated similarity of clades in *Shewanella* species.

| Clade | Described species included | No. of<br>species | G+C<br>content<br>(mol%)* | MLSA<br>concatenated<br>similarity (%) |
| --- | --- | --- | --- | --- |
| Algae | *S. algae*, *S. carassii*, *S. chilikensis* and *S. indica* | 4 | 53-54 | 94.8-96.6 |
| Amazonensis | *S. amazonensis* and *S. litorisediminis* | 2 | 54 | 91.2 |
| Aquimarina | *S. aquimarina*, *S. loihica* and *S. marisflavi* | 3 | 50-53 | 89.0-93.4 |
| Benthica | *S. benthica*, *S. psychrophila* and *S. violacea* | 3 | 47-49 | 90.6-94.6 |
| Colwelliana | *S. colwelliana* and *S. algidipiscicola* | 2 | 46-47 | 85.4 |
| Fodinae | *S. fodinae* and *S. dokdonensis* | 2 | 50-51 | 87.4 |
| Gaetbuli | *S. gaetbuli* and *S. aestuarii* | 2 | 43 | 84.3 |
| Hanedai | *S. hanedai* and *S. woodyi* | 2 | 44-46 | 87.0 |
| Japonica | *S. japonica*, *S. electrodiphila* and *S. olleyana* | 3 | 43 | 87.7-89.6 |
| Livingstonensis | *S. livingstonensis*, *S. algicola*, *S. arctica*, *S. basaltis*, *S. inventionis* and *S. vesiculosa* | 6 | 43-44 | 85.1-91.8 |
| Pealeana | *S. pealeana*, *S. fidelis*, *S. halifaxensis*, *S. kaireitica*, *S. marinintestina*, *S. piezotolerans*, *S. pneumatophori*, *S. sairae* and *S. schlegeliana* | 9 | 44-46 | 84.0-93.5 |
| Putrefaciens | *S. putrefaciens*, *S. baltica*, *S. decolorationis*, *S. glacialipiscicola*, *S. hafniensis*, *S. morhuae*, *S. oneidensis*, *S. profunda*, *S. seohaensis* and *S. xiamenensis* | 10 | 46-50 | 84.6-96.3 |

\*Calculated based on the concatenated sequences of six HKGs.

### Population genetics of *Shewanella* species in China

Eighty-six *Shewanella* strains isolated from diverse samples were involved in the analysis of sequences and phylogeny to evaluate the intraspecies relationships and investigate the distribution of *Shewanella* species in China. As shown in the concatenated phylogenetic tree (Fig. 4), eighty-six strains were divided into six compact clusters with high bootstrap support of 100 %. Each cluster was represented by a unique type *Shewanella* strain situated in Algae and Putrefaciens clades. In comparison with the concatenated phylogenetic tree, several unexpected locations were observed in the single HKG tree: strain 08MAS2647 in the *S. algae* cluster fell into the *S. chilikensis* cluster in *gyrA;* strains 08MAS2647, 11MAS2711, 11MAS2745 and 11MAS2746 in the *S. algae* cluster formed a subcluster next to the *S. carassii* cluster in *infB*; strains in *S. algae*; *S. carassii* and *S. chilikensis* clusters exhibited a close affiliation that could not be separated from each other in *recN*. Although some strains could be grouped into clusters properly in individual phylogenetic trees, clusters were supported with low bootstrap values, such as *S. algae* cluster in 16S rRNA, *gyrA* and *infB* genes as well as *S. seohaensis* cluster in *gyrA* and *gyrB* genes (Fig. S4). Hence, concatenated sequences derived from six HKGs exhibited good performance and robustness in identifying *Shewanella* strains. Since strains were defined as corresponding species in the concatenated phylogenetic tree, ranges of intraspecies and interspecies similarities for genes among the fifty-six validated *Shewanella* species were measured and shown in Fig. 5. The overlap between the intraspecies and interspecies similarities were observed among genes of 16S rRNA, *gyrA, infB, recN, rpoA* and *topA*. A small interval was detected in the *gyrB* gene with only 0.1 % variance. A notable gap was discovered in concatenated sequences. The minimum intraspecies similarity was found among *S. seohaensis* strains (97.8 %), and the maximum interspecies similarity existed between species *S. chilikensis* and *S. indica* (96.8 %), which differed by 1 % variation, corresponding to approximately 40 bp divergences.

**Figure 4.**
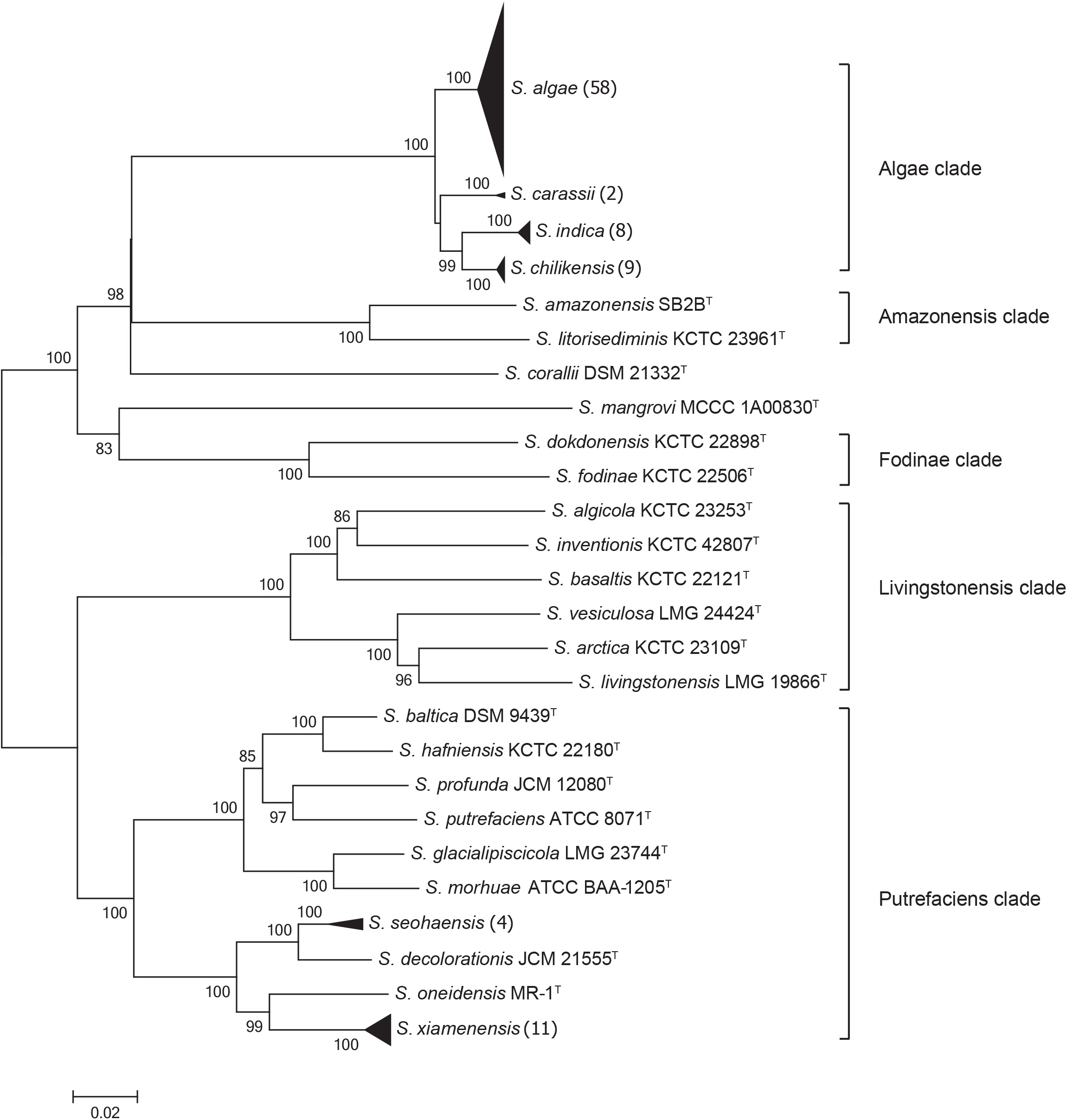
Phylogenetic tree reconstructed by the neighbor-joining method based on concatenated six gene sequences (*gyrA, gyrB, infB, recN, rpoA* and *topA*, 4191 bp) of eighty-six *Shewanella* tested strains and twenty-six related type strains. The strain number of tested strains for each compact cluster (black triangle) is shown in parentheses. The robustness of tree topologies was evaluated with 1000 bootstrap replications, and values greater than 70 % were shown at nodes of branches. The scale bar indicates substitutions per site.

**Figure 5.**
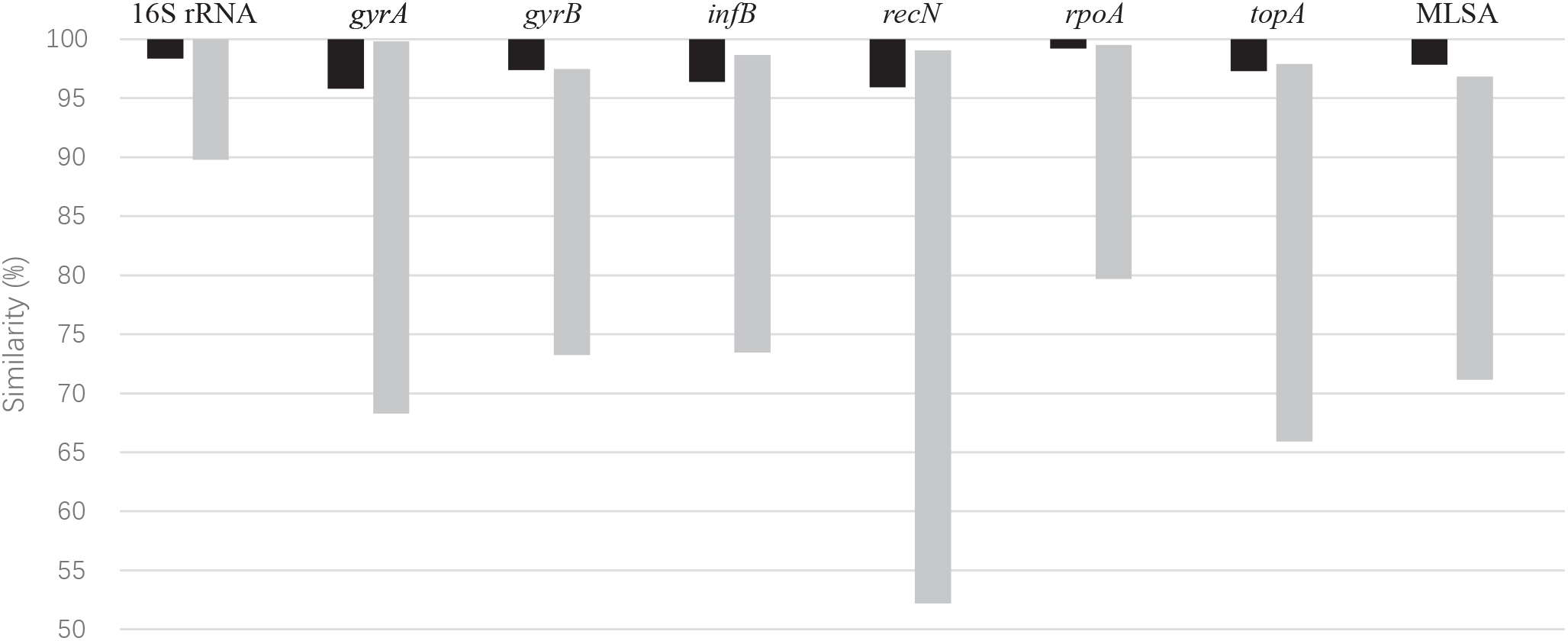
Intraspecies and interspecies similarities of 16S rRNA, six HKGs and MLSA for fifty-six validated *Shewanella* species. The ranges of similarity are displayed in black (intraspecies) and gray (interspecies).

Eighty-six *Shewanella* strains collected from China were subjected to define species via the MLSA approach. The most dominant *Shewanella* species was identified as *S. algae* (66.3 %), followed by *S. xiamenensis* (11.6 %), *S. chilikensis* (9.3 %), *S. indica* (8.1 %), *S. seohaensis* (3.5 %), and *S. carassii* (1.2 %). Except for the species *S. seohaensis*, which was only isolated from the environment, the remaining five species were relevant to clinical patients. It is noteworthy that species *S. algae, S. xiamenensis, S. chilikensis*, and *S. indica* were also discovered in food samples consisting of both marine products and cooked food for sale. Consequently, MLSA as a proper discrimination for *Shewanella* species played a significant role in public health and regular surveillance.

## DISCUSSION

In this study, the MLSA scheme, based on six HKGs (*gyrA, gyrB, infB recN, rpoA* and topA), was established for the first time to carry out efficient classification, reflect evolutionary relationships and delineate population biology in the genus *Shewanella*. Fifty-nine recognized type strains and eighty-six Chinese strains were investigated to explore the interspecies and intraspecies sequence diversity and phylogenetic topology in *Shewanella* species.

Previously, the 16S rRNA gene was applied as a traditional genetic marker among the genus *Shewanella* (17, 36, 37). However, the resolving power of the 16S rRNA gene was restricted with fewer parsimony informative sites and lower nucleotide diversity values. A narrow range of sequence variation was observed, and multiple pairs of *Shewanella* species shared greater than 99 % similarity. The latest proposed threshold of 98.65 % for 16S rRNA was insufficient to differentiate species in the genus *Shewanella* (38). Additionally, the existence of sequence variation among rrn operons would perplex the species definition and evolutionary analysis for taxa (39). Hence, protein-coding genes with a greater genetic resolution were utilized to determine the taxonomic position of *Shewanella* species.

Comparable analysis was performed among six HKGs (*gyrA, gyrB, infB, recN, rpoA* and *topA*). Unexpected classification of tested strains was discerned in the *gyrA* and *infB* genes for the high biological diversity among *S. algae* strains. The high interspecies similarities of those HKGs were generated, making them difficult to discern closely related species. The *gyrB* gene has always been used as a basic detection for novel *Shewanella* species identification (17–20). However, the criterion for *gyrB* analysis was not well established, and the boundary between interspecies and intraspecies similarities was inconspicuous. The *recN* gene was the most variable HKG with the greatest rates of parsimony informative sites and the widest spectrum of interspecies similarity. Although the *recN* gene was unsuccessful in making a distinction in the Algae clade, the effective discrimination was proven by high sequence substitution rates in the majority of species. The *rpoA* gene was more conserved than other HKGs with limited variable sites. None of the tested strains were phylogenetically located at unexpected positions, and only slight overlap was detected between intraspecies and interspecies ranges. The *topA* gene possessed a high genetic divergence next to the *recN* gene. The unstable taxonomic subtree with a low bootstrap value was discovered in the Colwelliana clade. The various evolutionary rates and inconsistencies of phylogenetic topology were discovered in those six loci. Therefore, the concatenated sequences with integrated and sufficient information should be taken into account to obtain the exact *Shewanella* species classification.

The concatenation of six HKGs demonstrated enough resolution power to discern *Shewanella* species in regard to variable sites, sequence divergences and independent branches. A notable gap between the ranges of interspecies and intraspecies similarities was favorable for defining the strains unambiguously at the species level, and 97.3 % MLSA similarity was proposed as a species threshold in the genus *Shewanella*. The neighbor-joining phylogenetic tree indicated that all validated species positioned at a distinct branch were clearly separated from closely related taxa. The stability of the phylogenetic tree was proven by bootstrap and topology analysis. The concatenated sequences phylogeny was supported by high bootstrap values among interspecies having a significant advantage over all individual genes. The phylogenetic tree grouped *Shewanella* strains into intraspecies clusters and taxonomic clades with almost 100 % bootstrap support. The use of the maximum-likelihood method had a slight impact on the tree topology. The reliability of the MLSA scheme was validated by comparison with genomic sequences. The identical phylogenies were constructed by concatenated sequences of six HKGs and core genes. A high correlation between the similarities of the MLSA and *is*DDH was discovered. Combined with the analysis of resolution, stability and reliability for nucleotide sequences and phylogenies, the MLSA approach of six HKGs (*gyrA, gyrB, infB, recN, rpoA* and *topA*) showed a significant performance for the precise classification of *Shewanella* species.

Under comprehensive analysis, the exceptional cases were only observed among two sets of recognized species, i.e., species *S. algae-S. haliotis-S. upenei* and *S. japonica-S. pacifica*. Based on molecular, genomic and phenotypic analyses, these five species were reclassified correctly, and the taxonomic structure of the *Shewanella* genus was refined. It is noteworthy that previous studies proposing those five novel species depended largely on the individual sequence analysis of 16S rRNA, experimental DDH and biochemical tests (7, 40–42). The high sequence similarities of 16S rRNA between their phylogenetic neighbors have already been observed, and the results of wet-lab DDH below 70 % were regarded as the gold standard for species classification (43). However, the experimental DDH was hard to reproduce completely by different laboratories; thus, the digital DDH based on the bacterial genomes was recommended in microbial systematics (23, 24). The phenotypic traits are inclined to be conservative among the *Shewanella* genus, and limited characteristics are suitable to discriminate *Shewanella* species. The deviation of biochemical results could be attributed to the different manual procedures and bacterial growth statuses. The phenotypic discrepancies in growth conditions and the carbon source utilization observed among species *S. japonica* and *S. pacifica* were also reported in the reclassification of species *S. affinis* and *S. colwelliana* (44). Therefore, the accurate molecular method of MLSA is considered a promising alternative tool for species identification and is superior to genomic analysis in terms of high efficiency and low cost.

In addition, the MLSA scheme provided a portable and robust system to reflect evolutionary relationships for the genus *Shewanella*. Twelve distinct phylogenetic clades were proposed with identical G+C contents and greater nucleotide similarity in concatenated sequences. The Chinese strains collected from clinical specimens and routine monitoring were located on Algae and Putrefaciens clades. These results indicated that species in monophyletic clades have a tendency to share a close genetic relationship, tracing back to common ancestry, and occupy similar geographical positions. These clades could be almost retrieved from individual HKG phylogenies, further elucidating the accurate and stable evolutionary structure in *Shewanella* taxon. Eight orphan species separated from all phylogenetic clades were defined. Attempts to involve the remaining species and identify the novel *Shewanella* species was conducive to exploring taxonomic positions for these species. In summary, the concatenated phylogeny provided significant insight into the evolutionary structure of the *Shewanella* genus for the first time.

Furthermore, it has been verified that *Shewanella* species, as marine pathogens, are associated with human diseases (12). Misidentifications to the species level were fairly common in clinical diagnoses due to the poor discernment system (45). In this study, eighty-six *Shewanella* strains collected from the environment, food and clinical samples in China were mainly defined as *S. algae*, followed by *S. xiamenensis, S. chilikensis, S. indica, S. seohaensis*, and *S. carassii* via the MLSA scheme. Five *Shewanella* species were verified to have connection with the clinic, including *S. algae, S. carassii, S. chilikensis, S. indica* and *S. xiamenensis*. It was likely that some *Shewanella* pathogens identified as *S. algae* in previous studies were believed to be *S. carassii, S. chilikensis*, and *S. indica* for their high 16S rRNA similarities. Apart from species *S. carassii*, four species were also frequently collected from marine products as well as cooked food for sale. It was reported that a common mechanism causing *Shewanella* infections was ascribed to the consumption of seafood or raw fish (12). Therefore, more attention is needed to reinforce continuous surveillance for the genus *Shewanella* by the MLSA approach in the processes of clinical diagnosis and food sales.

## ACKNOWLEDGMENTS

This work was supported by grants from the National Natural Science Foundation of China (31570134) and the National Sci-Tech Key Project (2018ZX10102001, 2018ZX10734404, 2018ZX10713001-002, 2018ZX10713003-002) from the Ministry of Health, China.

## Footnotes

The nucleotide sequences of six HKGs are deposited in GenBank nucleotide sequence database under accession numbers of *gyrA*: MH090144-MH090185; *gyrB*: MH090186-MH090202; *infB*: MH090203-MH090244; *recN*: MH090245-MH090286; *rpoA*: MH090287-MH090328; and *topA*: MH090329-MH090370

